# Challenges and opportunities for integrating genetic diversity monitoring in megadiverse countries: reflections from Mexico

**DOI:** 10.64898/2026.09.26.754592

**Authors:** Alicia Mastretta-Yanes, Libertad Arredondo Amezcua, Alexander Llanes-Quevedo, María Camila Latorre-Cárdenas, Juan Francisco Ornelas, Santiago Ramírez Barahona, Eugenia Zarza, Rossana Amaro-García, Francisca Acevedo Gasman, Valeria Alavez, Caroline Burgeff, Andrea Cruz-Angón, Rusby G. Contreras-Díaz, María Andrea Orjuela Restrepo, Oswaldo Oliveros-Galindo, Melissa Ríos-López, Diana María Rivera-Rodríguez, José de Jesús Sánchez-González, Sofía Suárez-Acuña, Jessica M. da Silva, Sean Hoban, Viktoria Köppä, Linda Laikre, Ana Wegier

## Abstract

Megadiverse countries concentrate most of the World’s biodiversity. These countries encompass different levels of research infrastructure, and nearly all of them have been historically committed to the Convention of Biological Diversity, often setting precedents and leading new approaches. Here, we ask whether the assessment of genetic diversity indicators provide novel insights for biodiversity management and conservation in one of those countries, Mexico. We evaluated the first national application of the Kunming-Montreal Global Biodiversity Framework (GBF), genetic diversity indicators in Mexico. The data covered 97 taxa across diverse ecosystem types and taxonomic groups, including crop wild relatives, to assess the capacity of genetic diversity indicators to provide novel insights, as well as their relevance for strengthening national conservation strategies. We calculated two indicators: the proportion of populations with an effective population size greater than 500 (Ne 500) and the proportion of populations maintained (PM). Data were sufficient to estimate the Ne 500 indicator for 47 taxa and the PM for 28 taxa, with mean indicator values of 0.217 and 0.936, respectively. Species’ geographic range was positively associated with Ne 500, whereas national endemicity had no significant effect. Comparisons with the IUCN Red List and Mexico’s national extinction-risk legislation revealed that genetic vulnerability occurs even among taxa not currently recognized as threatened. Our findings highlight a gap in legal recognition of genetic diversity protection that could be strengthened by incorporating population-level assessments focused on maintaining adaptive potential in the long-term. Additionally, the indicators allowed to highlight conservation concerns within crop wild relatives, though fully realizing their potential requires integrating the entire wild-to-domesticated complex. The Mexican experience highlights that the main challenge for scaling the monitoring of genetic diversity is not only generating new data, but integrating existing information into systems that strengthen collaboration and communication among researchers, local communities, and government institutions.

## INTRODUCTION

Genetic diversity is explicitly included in the Kunming-Montreal Global Biodiversity Framework (GBF) as part Goal A, which states: “[…] the genetic diversity within populations of wild and domesticated species, is maintained, safeguarding their adaptive potential” (CBD 2022a, 2022b). This marks a transition from biodiversity conservation primarily focused on species and ecosystems toward a more integrative approach that explicitly includes within-species variation as a measurable dimension of biodiversity that should be conserved (Hoban et al. 2024b; Mastretta-Yanes et al. 2024a). Following the first round of reporting since the adoption of the GBF, 13% of Parties reported the headline indicator on genetic diversity (A.4, “the proportion of populations within a species with an effective population size above 500”; CBD 2026; CBD2022b). Therefore, this is a crucial moment to reflect on the context, challenges and opportunities faced by the countries that reported on genetic diversity, as well as to examine whether their findings provide novel insights for how biodiversity is managed and conserved. Here, we focus on Mexico, a megadiverse country, where such a consideration is particularly necessary given its rich environmental heterogeneity, deep evolutionary history, complex biocultural systems and the resulting strong population structure and high levels of intraspecific diversity (Mastretta-Yanes et al. 2015).

Building on the recognition of genetic composition as a key attribute within the Essential Biodiversity Variables (EBVs) framework, genetic diversity has become a relevant dimension for global biodiversity monitoring systems (Laikre et al. 2020; Hoban et al. 2022; O’Brien et al. 2022). This shift was facilitated by the operationalisation of the genetic diversity indicators that are based on assessing population persistence and whether populations are large enough to retain genetic diversity (Hoban et al. 2020), which were adopted as complementary and headline indicators of Target 4 and Goal A of the GBF (CBD 2022a, 2022b). The complementary indicator focuses on population persistence, it reflects that when a population disappears, whatever genetic variation it contained is lost too, i.e. locally unique alleles. This metric, called “the proportion of maintained populations within species” indicator (PM indicator hereafter), is estimated as the number of populations that currently exist divided by the number of populations that originally existed. The headline indicator A.4 focuses on contemporary effective population size (Ne), a key population genetics metric that measures the rate at which populations lose genetic heterozygosity by genetic drift, thus losing adaptive potential (Allendorf et al. 2024). Importantly, the contemporary Ne reflects the capacity of the population to retain its current diversity going forward, regardless of the amount of diversity it currently has (Ryman et al. 2019). The indicator is estimated as the number of populations with Ne > 500 over the total number of populations for which there is available data (Ne 500 indicator). A threshold of Ne 500 is used because when Ne exceeds 500, evolutionary potential is expected to remain nearly stable, e.g., there is a very slow rate of heterozygosity decay (Hoban et al. 2024b). However, notice that allelic variation may go undetected if monitoring Ne alone (Allendorf et al. 2024). Therefore, the PM and Ne 500 indicators are most informative when considered jointly, and if they are interpreted considering fluctuations in census size (Nc; Hoban et al. 2026).

Taken together, the overarching goal of the Ne 500 and PM indicators is to enable the global assessment of whether species can maintain their adaptive potential, so that they can face environmental change and ongoing evolutionary pressures. They were developed aiming to balance scientific meaningfulness, practicality, and simplicity for countries from all over the world to report on them (Laikre et al. 2008, 2020; Hoban et al. 2024b). The central challenge was to develop simplified and standardised metrics that could reduce reliance on complex (and expensive) genomic analyses while retaining ecological interpretability and policy relevance (Laikre 2010; Hoban et al. 2024b). To balance this trade-off, the genetic diversity indicators rely on a pragmatic approach in which populations can be defined based on geographic or ecological variables. Similarly, the Ne of a population can be estimated from DNA-sequencing data or demographic census size (Nc) data using a Ne/Nc ratio (Hoban et al. 2024b; Mastretta-Yanes et al. 2024a). This allows the application of the indicators at large scales even in the absence of direct genomic data, keeping in mind that they should be considered complementary to DNA-based studies for fine-scale assessment (Hoban et al. 2024a).

A recent multinational effort, encompassing approximately 900 species across nine countries, demonstrated the feasibility of calculating at least one of these indicators for most taxa (∼83%) and revealed widespread and previously under-recognised loss of genetic diversity (Mastretta-Yanes et al. 2024a). Notably, the indicators pointed to loss of genetic diversity even in species categorized as Least Concern by the IUCN Red List, highlighting their complementary value to species-level assessments. This shows that for genetic diversity to be adequately conserved, it should be part of national assessment frameworks, particularly those embedded in legal and policy contexts at national levels. For instance, in Mexico, extinction-risk assessments are incorporated into formal policy instruments such as the NOM-059-SEMARNAT (NOM-059 hereafter), which consists of a list of species under extinction risk, and the method to generate it, that are protected by Mexican Law. The NOM-059 evaluates extinction risk based on species distribution, population size, habitat condition, and threats (SEMARNAT 2019). Although it incorporates elements of intrinsic biological vulnerability, its genetic component is not standardised through quantitative, comparable indicators such as the Ne 500 and PM indicators.

The application and interpretation of the PM and Ne500 indicators depend on several methodological decisions, including how populations are delimited, temporal baselines, and the use of DNA-based Ne estimates or proxies based on demographic and occurrence data, all of which can influence inferred trends (Laikre et al. 2008; Mastretta-Yanes et al. 2024a; Hoban et al. 2024a). Consequently, comparability across countries and taxa is not solely a technical issue but also an epistemological one, reflecting differences in how biodiversity is defined, measured, and monitored across socio-ecological contexts. Moreover, many megadiverse regions coincide with centres of origin and diversification of domesticated species, where the interaction between natural and human-mediated selection continues to shape genetic resources essential for global food security (Vavilov 1992; Bellon et al. 2017). This adds an additional layer of complexity, as conservation assessments must account not only for wild biodiversity but also for socio-ecological systems that sustain ongoing evolutionary processes in wild-to-domesticated species complexes (species composed by different evolutionary regimes, some of them influenced by humans; Alavez et al. 2026).

In this study, we examine the first national implementation of GBF genetic diversity indicators in Mexico, a megadiverse country and centre of origin for multiple domesticated species. Using approximately 100 species spanning diverse taxonomic groups and ecological systems, derived from a recent multinational assessment (Mastretta-Yanes et al. 2024), we examine whether these indicators uncover genetic vulnerability that is not captured in existing national frameworks. We further consider how these indicators can strengthen monitoring systems in Mexico, and discuss their potential relevance for other megadiverse countries.

## METHODS

Data used here was previously collected by Mastretta-Yanes et al. (2024a) as part of a multinational assessment including nine countries. Here, we focus only on the data collected for Mexico. Briefly, distinct poplations were defined using a variety of methods as described by Mastretta-Yanes et al. (2024b), which for the case of Mexico included geographic boundaries, ocurrence at different biogeographic zones or ecological conditions, distinct morphological traits, genetic clusters, and their combinations. Population size data for each population were then compiled as either: a) effective population sizes (Ne) estimated from genetic data by previous studies, b) point census counts (Nc) from studies or field personal observations, or c) Semi-quantitative data on Nc coming from local knowledge holders, expert opinion or field observations, that were standardized to discrete values: substantially >5000, slightly > 5000, just under 5000 and considerably fewer than 5000 individuals. Where Ne was unavailable, a conservative Ne/Nc conversion ratio of 0.1 was applied to estimate Ne from Nc data. The Ne 500 indicator was calculated as the proportion of populations with Ne > 500 relative to total extant populations with available data to estimate Ne. The PM indicator was calculated as the ratio of extant to total known populations (extant + extinct) and was omitted when the number of extinct populations was unknown.

The aim for Mexico was to assess around 100 taxa as a pilot to evaluate the feasibility of calculating the genetic diversity indicators for the national report to the GBF coordinated by the National Commission for the Understanding and Use of Biodiversity (CONABIO), a government agency linked to reporting to the GBF. Since Mexico is a megadiverse country with hundreds of thousands of described species, accounting for around 10-12% of the world’s species diversity (CONABIO, CONANP, PNUD 2020), it was recognized from the onset of the study that this diversity could not be truly represented with 100 taxa. Additionally, for a large proportion of the Mexican species, biological information is limited to occurrence distribution at best. Thus, rather than performing a random selection of all possible species in the country, the aim was to leverage the already on-going research efforts of the academic sector interested in genetic diversity. In Mexico, this community started to systematically publish genetic studies at the population or phylogeographic level of both domesticated and wild species in the late 2000s (Piñero et al. 2008; Mastretta-Yanes & Piñero Dalmau 2009; Mastretta-Yanes et al. 2018a) and has since then grown considerably. These researchers are either taxon experts or work closely with them, and have recently performed sampling trips so they were well positioned to provide both published or unpublished data to calculate the genetic diversity indicators. Researchers from this community were invited to participate in a pilot to estimate the genetic diversity indicators when the indicators were still under discussion in preparation to be presented at the COP15’ meeting. The invitation was done in October-November 2022 via email to known active researchers on population genetics or phylogeography in Mexico, further amplifying the invitation within their institutions. The final Mexican team was composed of nine core people, including researchers and students from the academic sector, as well as a former staff member from CONABIO. Approximately 25 additional researchers, students, and CONABIO staff participated in workshop discussions or facilitated access to data.

A preliminary list of species or subspecies was compiled based on researcher expertise and anticipated data availability. Taxa were included when participating researchers had either previously conducted genetic studies on them or could have sufficient field knowledge to provide the information required for the assessment. From this list, the final list of taxa were chosen to: (1) represent the main taxonomic groups of terrestrial animals (amphibians, birds, fishes, invertebrates, mammals, reptiles) and plants (angiosperms, bryophytes, gymnosperms, pteridophytes), including crop wild relatives (CWR), i.e. taxa closely related to crops, or that the taxon itself is part of wild-to-domesticated complex of a crop; (2) to include endemic and non-endemic taxa, both widespread and range-restricted; and (3) to represent the main terrestrial biomes of Mexico.

Once the species list was defined, the core team worked closely with members of the other eight countries from Mastretta-Yanes et al. (2024a)’s to co-design and include the Mexican context into the overall guidelines (Mastretta-Yanes et al. 2024b) and shared questionnaire for data captured (developed using KoboToolBox, https://www.kobotoolbox.org/). Within the Mexican team, training and discussion sessions were undertaken online, including members of the core team, as well as undergraduate and postgraduate students. Besides the core group performing the assessments, taxon experts were reached to discuss some specific cases. Data sources included the scientific literature, CONABIO’s biodiversity information system accessed through Enciclovida (https://enciclovida.mx/), unpublished data including fieldwork notes and taxon expert opinion. Where uncertainty existed (e.g., in defining number of populations), taxa were assessed under multiple scenarios. All metadata was captured during this process, except for the NOM-059 extinction risk, and flagging which species are CWR, which were added during the preparation of this study.

Prior to the automatic data quality tests of Mastretta-Yanes et al. (2024a), each of the data entries for Mexican species was validated by a team member different from the person who submitted it. Three assessments did not pass the quality tests and were discarded. The resulting dataset was then downloaded as a.csv file and processed in R v. 4.4.3 using functions available at https://github.com/CCGenetics/Ginko-Rfun for data cleaning and processing. Plots and analyses specific for this study were done using custom functions available at XXXX (available upon acceptance). We used a generalized linear mixed model with a beta distribution to examine the effects of species geographic range and national endemicity on the Ne 500 indicator variable using the package glmmTMB v1.1.14 (Brooks et al. 2017). Additionally, we performed Kruskal-Wallis rank sum tests to non-parametrically assess the effects of species geographic range and national endemicity on the Ne 500 indicator variable.

## RESULTS

The assessments were done for a total of 97 taxa (species or subspecies, Table S1), including 44 non-endemic (19 animals and 25 plants) and 53 endemic (7 animals and 46 plants, Figure 2A). More plants (62: 3 pteridophytes, 8 gymnosperms, 51 angiosperms) than animals (36: 1 invertebrate, 1 fish, 2 amphibians, 9 reptiles, 17 birds, 5 mammals) were included. The bias towards plants occurred because 19 CWR were included, considering 10 cotton taxa (*Gossypium* spp.), five teosintes (wild relatives of maize, Zea spp.), one papaya (*Carica papaya*), one pumpkin (*Cucurbita argyrosperma ssp. sororia*), and two edible cacti (*Stenocereus* spp.), indicated by dots in Figure 3.

A total of 98 assessments were done where it was attempted to estimate the Ne 500 or PM indicators based on available data (Figure 1). The discrepancy between the number of taxa and assessments emerged because one taxon, *Alouatta palliata ssp*. mexicana (Mexican howler monkey), was assessed more than once. This was needed because despite that one study found three genetic clusters (Solórzano-García et al. 2021) and other a single cluster (Melo-Carrillo et al. 2020), experts argued that a greater effort is required to include more sampling sites, standardize the methods of using microsatellite markers, and consider possible cases of hybridization. Thus, *A. palliata* ssp. *mexicana* was evaluated as having either three populations or one.

**Figure 1.**
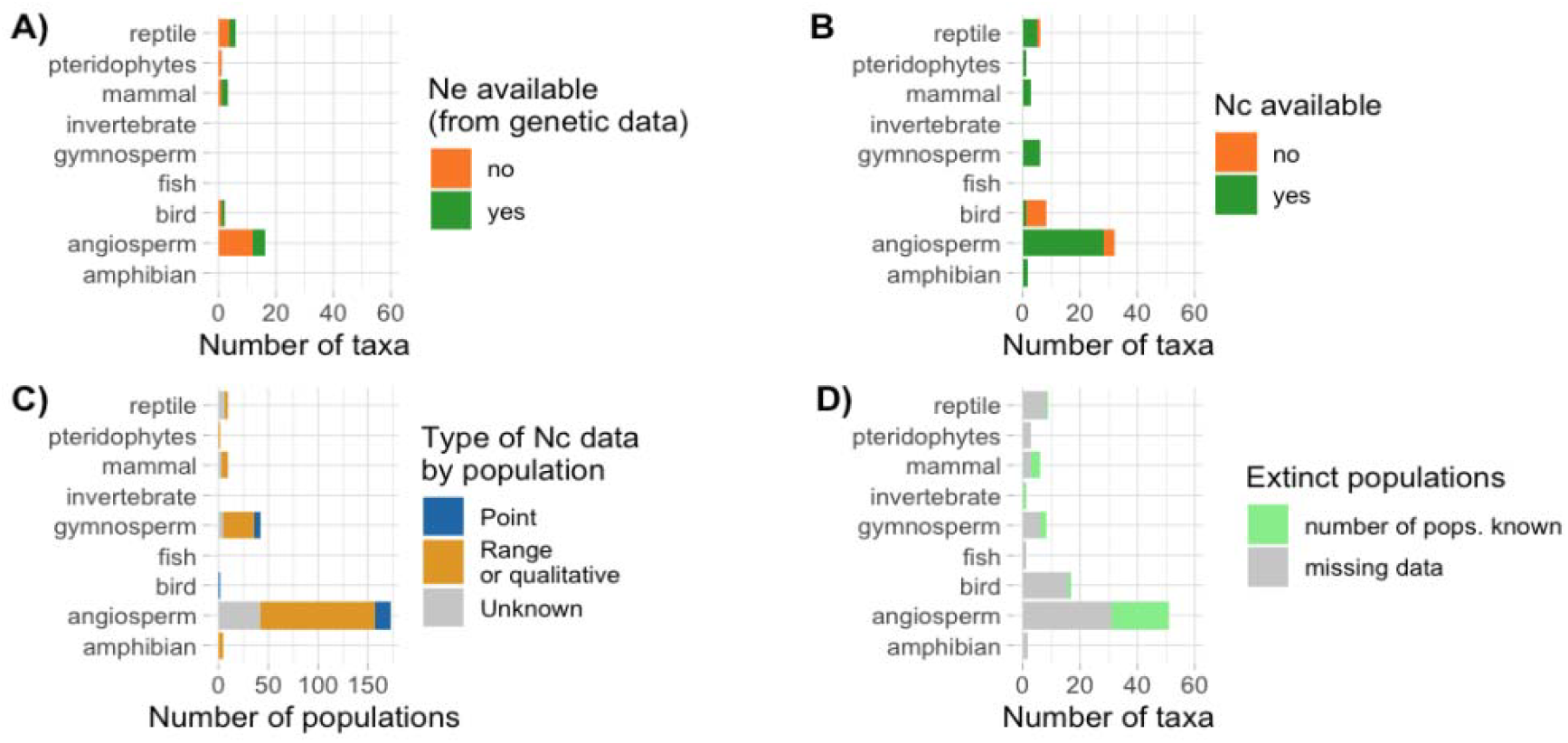
Data availability for the Ne 500 indicator (A-C) and the PM indicator (D). (A) Total number of taxa with effective population size (Ne) data estimated from DNA-based (genetic) data for individual populations. (B) Total number of taxa with census population size (Nc) data available for individual populations. (C) For taxa where Nc data was available, “point” data represents count estimates or point approximate values (e.g., capture–recapture study found 3120 individuals), and “range or qualitative” is more generic estimates of census size, either represented by quantitative ranges or qualitative descriptions of population size (e.g., “a few hundreds” or “>5000 by much”. (D) Total number of taxa with and without data on the number of extinct populations within a taxon, as needed for the PM indicator.

There was available data to compute the Ne 500 indicator (Figure 1A-C) in 47 (48%) of the taxa. Considering only these taxa, the indicator mean value is 0.217 (Figure 3A). That means that 21.7% of the analysed populations are large enough to prevent the loss of genetic diversity. For the PM indicator, there was available data (Figure 1D) to compute it in 28 taxa (28%), with a mean indicator value of 0.936 (Figure 3B). That means that in the analysed species 93.6% of their populations still exist. It was possible to estimate both indicators together only in 17 taxa (Figure 3A).

When examining indicator values across taxa, the Ne 500 indicator showed consistently lower values, with relatively few populations meeting the Ne ≥ 500 threshold (i.e. loss of *within* population diversity). In contrast, the PM indicator values were generally higher and more frequently reached the maximum value (PM = 1), indicating that for many taxa, no documented population loss was detected (i.e. maintenance of *between* populations diversity). This pattern holds across endemic and non-endemic taxa. However, the generalized linear mixed model revealed a significant positive effect of species geographic range on the Ne 500 indicator variable (p = 0.033), while national endemicity had a non-significant negative effect (p = 0.139, Figure 2B-D). The results suggest that more widespread species tend to have higher Ne 500 indicator values as compared to range-restricted species. These findings were corroborated by Kruskal-Wallis rank sum tests, which showed a significant effect of species geographic range (p < 0.001) and a non-significant effect of national endemicity (p = 0.063) on the Ne 500 indicator variable.

**Figure 2.**
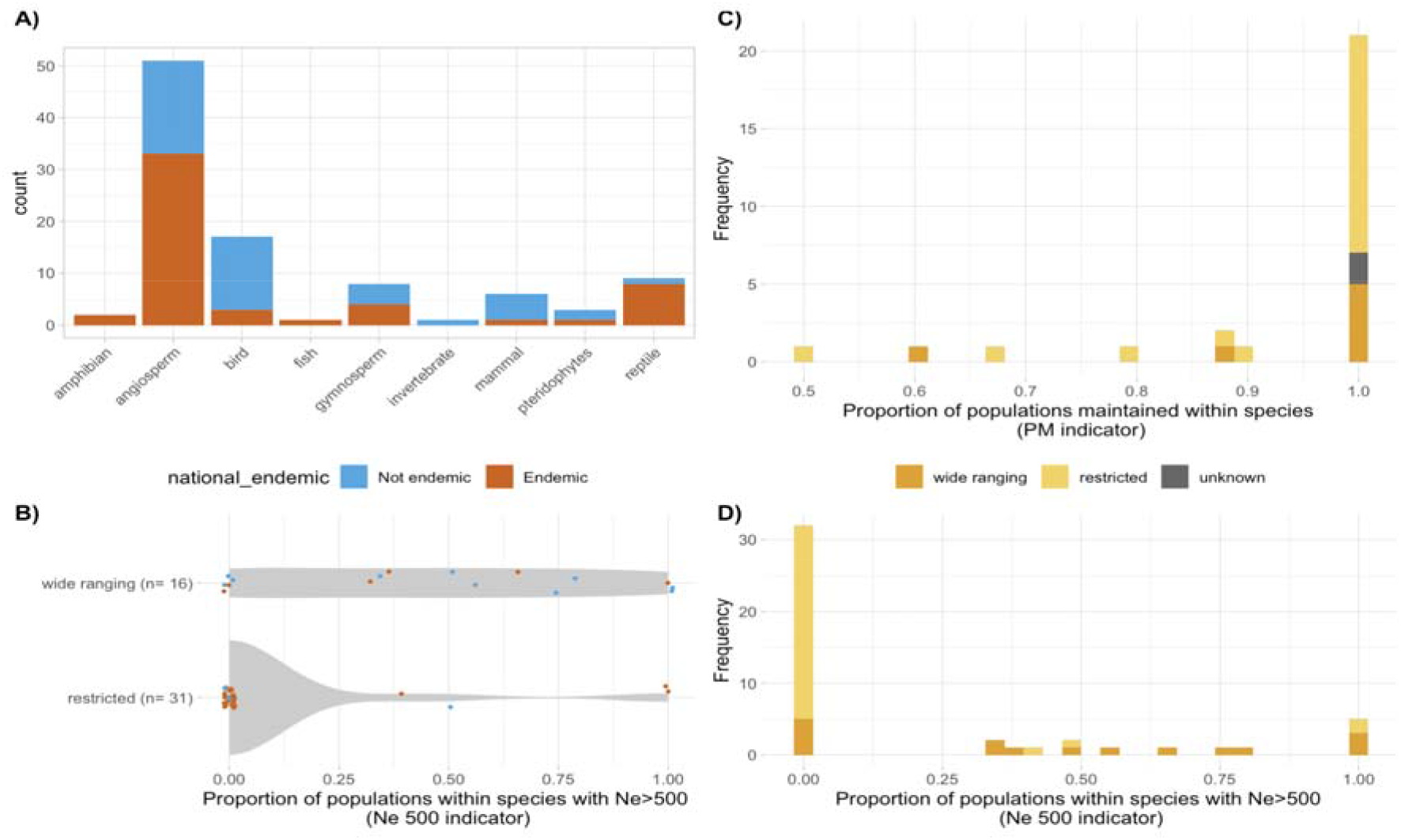
Assessed Taxonomic groups and indicator values by endemicity and range type. (A) Number of taxa assessed by taxonomic group and whether they are endemic to Mexico. (B) Violin plots showing the range in indicator values across species range types, for the Ne >500 indicator. Each dot represents the indicator value for a single assessment, coloured by endemicity, with the sample size, *n*, for each range type. (C-D) Frequency barplots for the PM and Ne 500 indicators, respectively, coloured by range type.

Regarding the extinction risk of the assessed species, the number of species by category are distributed in similar numbers across both the Global Red List from IUCN and the NOM-059, i.e. the estimation of the genetic diversity indicators for Mexico included from “none known” to the highest levels of threat (Figure 4A). The majority of the threatened (CR, EN, VU) according to the IUCN Red List are also listed in the NOM-059, and most species not assessed by the NOM-059 are assessed as least concern (LC) by the Red List (n=9). Interestingly, some species present in the NOM-059 have not been evaluated by the IUCN Red List (n=5), and there is an overlap between taxa not assessed by neither list (n=7, Figure 4A). Taxa categorized with some level of threat by either the Red List or the NOM-059 tended to show low values for the Ne 500 indicator (Figure 4 B-C). The “least concern” (LC) category of the Red List showed the highest spread of genetic diversity indicators values. This is similar to the pattern found for the other eight countries of Mastretta-Yanes et al (2024). This pattern is also reflected in the NOM-059 for the non-evaluated species (Figure 4C), with most of them (14/21, i.e. 66%) with a Ne 500 indicator value ≤ 0.5. That means that there is a high proportion of taxa not currently included in the NOM-059 where at least half of their populations, or even all, are losing genetic diversity because they are too small.

## DISCUSSION

### Megadiversity as a structural condition for interpreting genetic diversity indicators

Megadiverse countries tend to be more committed in their reports to the CBD than non-megadiverse countries (Maney et al. 2024). Congruent with this, Mexico was one of the 13% of Parties that reported on the genetic diversity indicator (headline A4), using the data discussed here. The national average for this indicator (indicator value 0.217) is based on 47 taxa that had population-level Nc or Ne to allow the calculation of the Ne 500 indicator. This is similar to the number of taxa that other countries reported on. Regarding data availability, the patterns observed in Mexico also align with findings from other countries participating in the multinational assessment (Mastretta-Yanes et al. 2024a). As in other countries, approximately half (47%) of the assessed species had sufficient data to estimate the Ne 500 indicator, while fewer (28%) provided adequate data for PM indicator calculations. Notably, Ne estimated directly from genetic data was rarely available; instead, Ne was most often estimated from Nc data, derived either from point estimates or expert approximations of the number of mature individuals, including rough estimates (e.g., “much more than 5,000). On one hand, this shows that there is an opportunity for genetic studies to contribute to national monitoring efforts by including the estimation of Ne. On the other hand, the availability of Nc data shows that involving non-geneticist experts, as done for instance during NOM-059 and Red List assessments, would yield useful data (Khorngton et al. 2026).

Altogether, what Mexico has demonstrated is that it is possible to establish an initial national baseline from heterogeneous existing information, even in a megadiverse country and despite taxonomic and data limitations. Of course, the taxa list should be considered an informative first step, not fully representative of the Mexican biodiversity; it should be subsequently extended targeting different biomes, taxonomic groups and national priorities. Immediate concerns for Mexico include aquatic ecosystems, a biome that remains significantly underrepresented in current monitoring efforts globally. Also, as a centre of origin, domestication and diversification for numerous crops (Harlan 1971; Casas et al. 1997), Mexico provides an exceptional setting to evaluate CWR, as will be discussed later. However, since monitoring in any megadiverse country is particularly challenging (Moussy et al. 2022), it is likely that expanding the list of taxa assessed to hundreds would not fully account for its vast diversity. Most importantly, monitoring requires measurements to be repeated consistently enough to detect change, so there is a tradeoff between monitoring more taxa and repeating assessments through time.

Even assessing relatively few Mexican taxa for this baseline, the genetic indicator values revealed two interesting patterns. First, that taxa with restricted geographic ranges consistently show lower indicator values compared to species with broader more widespread distributions (Figure 2); and second, that even taxa classified as Least Concern by the IUCN or less level of risk by the NOM-059 exhibit appreciable loss of within-population genetic diversity (Figure 4B,C). The first pattern carries particular significance for megadiverse countries. The exceptional species diversity characterizing these regions is substantially explained by an overrepresentation of geographically restricted species, which tend to therefore be endemic (Jetz et al. 2004). However, an important distinction emerges between “range type” and “endemism” per se as predictors of indicator values. Although endemism is fundamentally a biogeographic concept, its practical application frequently conflates it with political boundaries of varying scales (Daru et al. 2020). In a large country like Mexico, for example, endemic species may possess extensive distributions spanning thousands of kilometres (e.g. the iguana *Ctenosaura pectinata*, assessed here; Zarza et al. 2008). Range type, by contrast, more directly captures the ecological and biogeographic factors that structure intraspecific genetic variation.

In Mexico, the high number of species with restricted distributions is explained by the presence of tropical mountains, which create naturally fragmented landscapes (Koleff et al. 2008). Importantly, these mountainous regions support populations capable of persisting through climate fluctuations and, critically, tend to harbour strong signals of population genetic differentiation (Ramírez-Barahona & Eguiarte 2013; Mastretta-Yanes et al. 2015, 2018b). This geographic structure has profound evolutionary implications: while the PM and Ne 500 indicators remain numerically comparable across taxa and countries, the genetic distinctiveness lost in megadiverse tropical regions likely involves deeper losses than what occurs at higher latitudes, where Pleistocene fluctuations in general generated more homogeneous population structures with reduced accumulated differentiation among populations (Hewitt 2004; McDevitt et al. 2021; de Jong et al. 2023). For example, among the species assessed here are *Juniperus blancoi*, a conifer in which populations (some of them recognized as subspecies) from central and north Mexico diverged around 6 million years ago (Moreno-Letelier et al. 2014); as well as several species distributed in cloud forests, with divergence times among populations at different sides of the Tehuantepec Isthmus spanning from thousands to millions of years (e.g., the hummingbird *Lampornis amethystinus* and the cactus *Rhipsalis baccifera*, respectively; Ornelas et al. 2013). Thus, given that high levels of population differentiation are common, for Mexico it would be particularly important to assess both the PM and Ne 500 indicators together as a minimum priority for biodiversity monitoring.

Regarding the second pattern, the fact that even non-endangered taxa have low indicator values (Figure 4), means that the extinction-path starts with the quiet decrease of population size at local levels, followed by local population extinction. Our results suggest that this loss of genetic diversity goes unnoticed by the assessment of the Global Red List of the IUCN or the NOM-059, showing that monitoring genetic diversity provides an additional insight beyond what is covered by current biodiversity assessments. Thus, in the context of environmental law, our findings point to a gap in the legal recognition and protection of genetic diversity within the Mexican legislation and regulations, which is likely the same case elsewhere.

### Genetic diversity indicators reveal a vulnerability not addressed by the current Mexican legislation

One characteristic of environmental policy in Mexico, as in many other countries, is that its current legal and institutional framework has evolved through the interaction between national priorities and international commitments. International agreements often establish broad conservation goals that are subsequently incorporated into national legislation and progressively strengthened through national scientific knowledge, institutional experience and policy development (Brand & Görg 2003). The NOM-059 illustrates this process. The first official Mexican standard listing species at risk was NOM-059-ECOL-1994, published in 1994, after Mexico’s ratification of the Convention on Biological Diversity. Since then, it has been continuously refined through national expertise, specialist participation and periodic technical assessments (Pavón et al. 2023). Today, listing a species under NOM-059 formally recognizes it as being at risk under Mexican law. In practice, this status informs environmental impact assessments, land-use planning, permitting decisions, and conservation and recovery measures, while providing a legal basis for implementing species-specific protection actions. However, our results reveal an important limitation of NOM-059: it focuses on assessing species already considered likely to be at risk. Consequently, many species that have never entered the assessment process may already be experiencing reductions in Ne sufficient to compromise their long-term evolutionary potential, as shown in Figure 4C. Likewise, species currently included in the NOM-059 may face greater conservation risks than previously recognized, because the NOM-059 was not designed to support systematic monitoring of genetic diversity through repeated assessments over time using a consistent methodology, making it difficult to compare changes in genetic status across assessment periods. In this context, the adoption of the genetic diversity indicators under the GBF represents an opportunity to complement existing conservation assessments at national level, by both extending the species to be assessed and including the genetic diversity indicators as part NOM-059 criteria.

Beyond adjustments to the NOM-059, there are other opportunities for recognizing genetic diversity in the Mexican environmental law. The NOM-059 is a legally binding technical standard issued under the broader legislative framework of the Ley General del Equilibrio Ecológico y la Protección al Ambiente (LGEEPA, General Law on Ecological Equilibrium and Environmental Protection), the federal law that establishes the general legal principles and obligations for environmental protection and biodiversity conservation. One of the primary objectives of the LGEEPA is to establish the legal basis for guaranteeing people’s right to live in a healthy environment, which is one of the rights recognized in the Mexican Constitution (Article 4, fifth paragraph). Genetic diversity is related to maintaining a healthy environment, because it underpins the capacity of species to adapt to environmental change, thereby contributing to the resilience and long-term functioning of ecosystems (Des Roches et al. 2018). Preserving the adaptive potential of species is therefore not only a biodiversity conservation objective, but also an essential component of maintaining a healthy environment which should be of direct relevance for LGEEPA (Mastretta-Yanes 2024). Nevertheless, the only explicit mention of genetic diversity within LGEEPA is on its article 45th, which enlists “safeguarding genetic diversity of wild species” as one of the objectives of establishing National Protected Areas. Beyond its role in justifying protected areas, genetic diversity could be explicitly recognized in the broader biodiversity provisions of the LGEEPA, particularly Article 79, which sets out the guiding criteria for the conservation and sustainable use of biodiversity.

Similarly to its relationship with the right to a healthy environment, genetic diversity is also relevant to the right to “adequate food that is nutritious, sufficient, and of good quality”. This constitutional right (Article 4, third paragraph) is further developed in the Ley General de la Alimentación Adecuada y Sostenible (LGAAS, General Law on Adequate and Sustainable Food), which emphasizes that the right to food encompasses sustainability, as well as biocultural richness and the relationship between food and culture. Genetic diversity contributes to these objectives by supporting more sustainable and nutritious agriculture, for instance through breeding programs that leverage traits from crop wild relatives to improve resistance to pathogens (Śliwka & Zimnoch-Guzowska 2013), by providing essential nutrients found in traditional crop varieties (Medina-Lozano & Díaz 2021), and by enabling the emergence and maintenance of locally adapted varieties of cultural significance (Mastretta-Yanes 2024). In a centre of origin and domestication such as Mexico, this inevitably drives the attention to taxa whose evolution occurs across wild–managed– domesticated gradients, opening the question of how to conserve and monitor their genetic diversity.

### Towards monitoring genetic diversity in wild-to-domesticated species complexes

This study was conducted in parallel with the development of the conceptual and methodological foundations for implementing the genetic diversity indicators of the GBF (Hoban et al. 2023; Mastretta-Yanes et al. 2024a, 2024b). In that context, the wild-to-domesticated species complex represented an additional layer of complexity, therefore we decided to start tackling this issue by focusing on CWR only, i.e. on wild species related to crops or only on wild populations for those species that occur in a wild-to-domesticated complex. This provided an opportunity to evaluate the performance of genetic diversity indicators in species that, because they are crop wild relatives or form part of wild–to-domesticated complexes, represent a priority component of Mexico’s biodiversity (Mastretta-Yanes et al. 2018a; Contreras-Toledo et al. 2019). Thus, in this study 19 of the 97 taxa analysed (19.6%) are CWR, including species of cotton, teosinte (wild relatives of maize), papaya and edible cacti (highlighted by black dots in Figure 3). In Mexico, these taxonomic groups are particularly relevant not only because of their contribution to food security and sovereignty (Goettsch et al. 2021), but also because they are embedded within a shared evolutionary and biocultural history in which biological and cultural diversity have co-evolved (Casas et al. 2007; Casas & Parra 2016).

**Figure 3.**
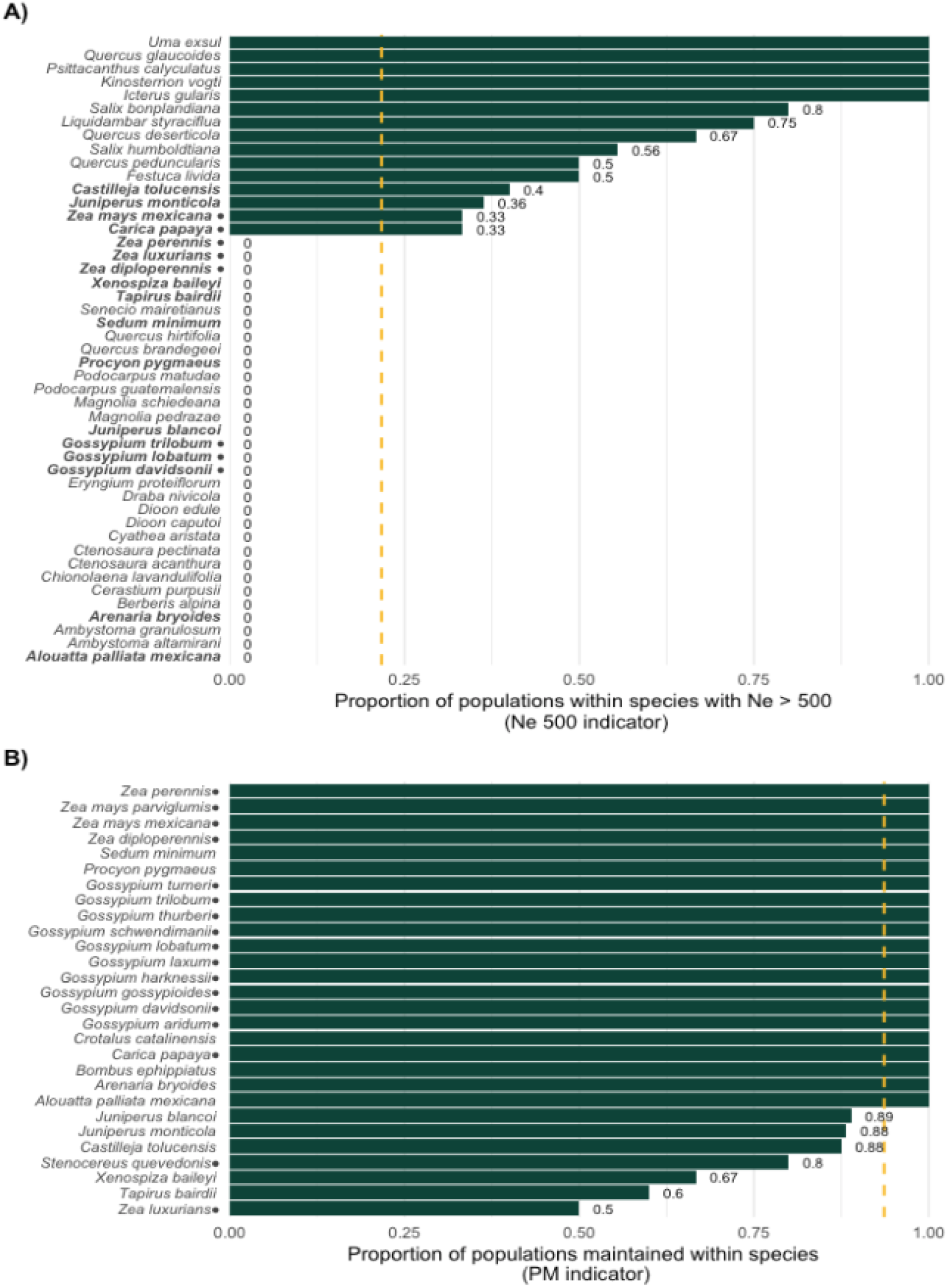
Genetic diversity indicators for each taxon for which the Ne 500 indicator (A) and the PM indicator (B) could be estimated. Taxa are ordered by indicator value. Taxa with an indicator value of 1 are indicated by full width bars. For the rest of the cases the individual indicator value is shown at the right of each bar. The dashed line shows the average across taxa for each indicator. Taxa for which it was possible to calculate both indicators are shown in bold in (A). Crop wild relatives are marked with a dot (⍰).

**Figure 4.**
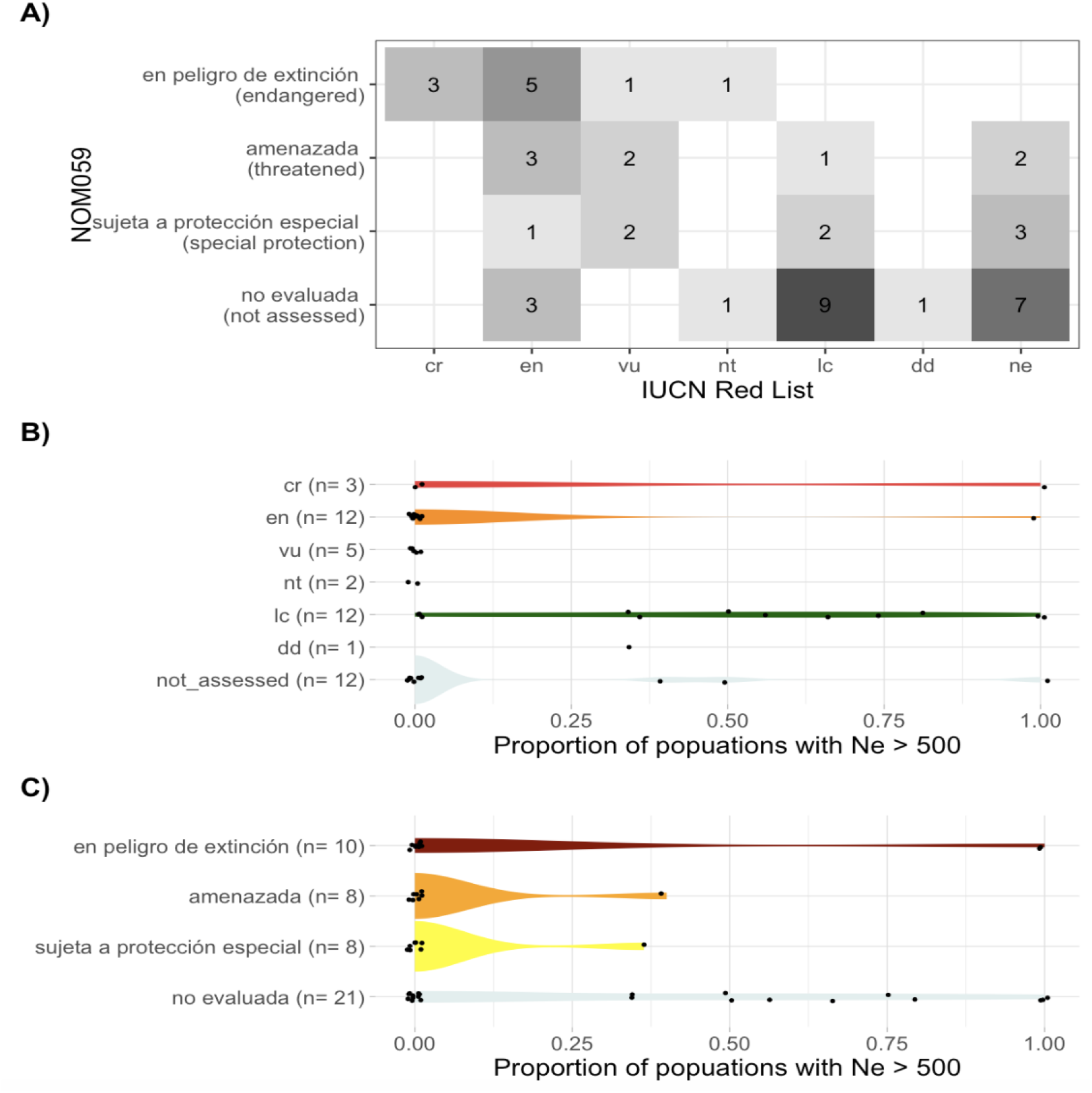
Genetic diversity indicators by extinction risk according to the Mexican NOM-059 and the Global Red List from IUCN. (A) heatmap of the number of species in each category for each of the lists. Each category of both evaluation methods is represented by at least one taxon. Abbreviations for the IUCN correspond to: cr (Critically Endangered), en (Endangered), vu (Vulnerable), nt (Near Threatened), lc (Least Concern), d (Data Deficient), and ne (Not Assessed). (B-C) Violin plots illustrating the spread in the Ne 500 indicator for the IUCN Red List and the NOM-059, respectively. Each dot represents the indicator value for a single assessment.

Our results demonstrate the ability of the indicators to identify conservation concerns for genetic diversity within CWR. The teosinte *Zea mays* subsp. *mexicana*, for example, retains all its populations (PM = 1), yet only approximately one-third have effective population sizes above the Ne = 500 threshold, indicating that losses of adaptive potential may precede population disappearance. A similar pattern was observed in the wild populations of *Carica papaya*, whereas the perennial teosintes *Zea diploperennis, Z. perennis*, and the wild cottons *Gossypium trilobum, G. lobatum* and *G. davidsonii* showed Ne 500 indicators values of zero, indicating that none of their populations maintain the effective population size considered necessary to preserve long-term adaptive potential. Likewise, the teosinte *Zea luxurians* and the cacti *Stenocereus quevedonis* already exhibit population loss (PM = 0.5 and 0.8, respectively), and the remaining populations of *Z. luxurians* also fall below the Ne = 500 threshold. Due to their importance for food security, certain CWR have received considerably funding for *in situ* studies and remain a focus of ongoing research efforts. For instance, populations from the perennial teosintes have been visited over the last two years by members of this team. This opens up the possibility of moving the genetic diversity indicators from baseline to monitoring.

We showed that the indicators can detect signals of genetic erosion even when only the wild component is evaluated. However, as recognised in evolutionary studies and conservation planning, excluding components of the wild-to-domesticated complex may lead to incomplete interpretations of evolutionary processes and, consequently, to incomplete conservation strategies (Cronin et al. 2020; Moreno-Letelier et al. 2020; Alavez et al. 2026). Although conceptual frameworks for addressing these continua in plants have recently been proposed (Alavez et al. 2026), their implementation within the context of the genetic diversity indicators remains a major methodological challenge. In the future, incorporating domesticated populations and other components of wild-to-domesticated species complexes may modify the assessment of genetic diversity in species such as *Carica papaya*, where modern and traditional cultivars (e.g., domesticated), feral and wild populations co-exist under the same taxonomic name, with gene-flow among them (Ruiz-Gil et al. 2023).

Our assessment also identified other challenges particularly associated with CWR. In Mexico, information on these species is distributed across research institutions and government agencies with objectives ranging from conservation biology and evolutionary research (e.g., Universities) to genetic improvement, agricultural, livestock and forestry production (e.g., Ministry of Agriculture and Rural Development, SADER), as well as the management and conservation of biocultural resources by Indigenous Peoples and local communities (e.g., Ministry of Environment and Natural Resources, SEMARNAT and the National Institute for Indigenous People, INPI). This reflects a global trend separating wild *vs* agriculture research (Jago et al. 2024). Consequently, future methodological approaches should be capable of both integrating information from disparate sources, and incorporating the multiple evolutionary trajectories that may coexist within a single species from a wild-to-domesticated complex. Another challenge associated with CWR is that, for some domesticated species, their wild relatives are not known with certainty; in such cases, the entire genus could need to be assessed until it is determined which species are part of the domesticated species’ gene pool.

Beyond their role in international biodiversity reporting, genetic diversity indicators can allow assessing whether the evolutionary and biocultural processes that generate and maintain biodiversity remain functional. Their ability to do so, however, depends on whether populations are defined in ways that adequately capture the evolutionary trajectories of the species being assessed. For instance, in wild-to-domesticated complexes of plants, genetic diversity is distributed across a continuum that may include wild, co-evolved, incipiently domesticated, semi-domesticated, fully domesticated, landrace, modern cultivar, feral and volunteer populations (Alavez et al. 2026). In this context, the explicit recognition within the GBF that genetic diversity should be monitored and conserved in both wild and domesticated species (CBD 2022a, 2022b) represents an important step towards incorporating biocultural evolutionary processes into biodiversity monitoring. Fully realising the potential of the genetic diversity indicators, however, will require methodological frameworks that explicitly incorporate wild-to-domesticated species complexes while integrating genetic evidence with ecological, historical and biocultural knowledge, together with the knowledge and management practices of farmers, Indigenous Peoples and Local Communities (IPLC). Expanding genetic indicators to these systems offers an opportunity to understand how well food production systems are positioned to face rapidly changing conditions.

### Incorporating genetic diversity into conservation requires transversal communication

This first national assessment demonstrates that Mexico already possesses many of the capacities required to implement genetic diversity indicators. The availability of biological information, institutional infrastructure, and scientific expertise made it possible to assess an initial set of taxa and, importantly, generated interest within the national research community in applying these metrics more broadly. For example, contemporary Ne started to be estimated from genomic data by new studies, involving students that attended seminars on the genetic diversity indicators undertaken by the Mexican team (Truchot-Taillefer et al. 2026; Martínez-Velasco et al. 2026). Thus, the main challenge identified by this study appears to be less the generation of new information than the integration of existing information into a workflow that can strengthen confidence in its management, use, and continuous update involving the academic and government sectors, but also indigenous peoples and local communities.

Academic knowledge is disseminated primarily through peer-reviewed publications, often in English and behind paywalls, and through postgraduate theses, following processes that can take several years. Although these outputs generate fundamental knowledge, they are rarely translated into formats that can directly inform governmental decision-making or biodiversity management (Morcillo et al. 2026). At the same time, local communities living alongside biodiversity cannot postpone management decisions affecting genetic diversity until scientific studies have been completed and their results published. Their decisions are thus grounded in knowledge accumulated over generations, cultural values, local experience, immediate needs, and, where available, scientific knowledge that has reached them before a decision needs to be made (Wheeler & Root-Bernstein 2020). Supporting local dialogue processes is therefore also important for identifying and assessing the information that people in direct contact with the populations being evaluated can provide and, together with other actors, interpret the indicators. Such dialogue can help identify threats and sustain the evolutionary processes that underpin biodiversity.

Because evolutionary processes are maintained at the population level, the conservation of genetic diversity must also take place at the population level. IPLC, protected-area managers, and citizen-science networks are often particularly well positioned to detect changes in populations over time. This is supported by this dataset, for instance 68% of the Nc data corresponded to Nc approximate ranges (Fig. 1C), obtained based on personal field observations of academics, park rangers or interviews with local knowledge holders. Through standardized monitoring protocols, these actors could continuously provide valuable information, strengthen awareness of the importance of conserving genetic diversity at local and regional scales, and implement management actions whose outcomes could be assessed within the same monitoring framework. In turn, an effective flow of information would allow local observations, and emerging conservation needs to reach government agencies and research groups able to address remaining knowledge gaps. Communication, therefore, is not simply an additional component of implementation; it is the process through which knowledge is exchanged, integrated, and mobilized, enabling adaptive monitoring and the progressive improvement of conservation actions.

Open-access biodiversity information systems could provide an important means of connecting these actors. In this study, Enciclovida (CONABIO’s data portal to the Mexican Biodiversity Information System, https://enciclovida.mx) enabled us to integrate and visualize observations from scientific collections, citizen science, explore the distribution and proximity of records, support the verification of localities to define populations and reach local knowledge holders or experts for population-level census size data. Species reports, which are often part of biodiversity information systems (CONABIO 2017), could also incorporate population-level data useful to systematically monitoring genetic diversity. Depositing, updating, and mobilizing information does not alter the conservation status of a species or population, but systematic and accessible information would strengthen the capacity to detect changes, respond in a timely manner, and conduct assessments at the national scale.

In other words, in practice, information systems already exist that can integrate data from multiple sources that could be useful to estimate the genetic diversity indicators. This lesson learnt from Mexico is relevant to other megadiverse countries, because the majority operate well-established digital public infrastructures (UNDP 2026) hosting national biodiversity information systems that centralize biological data through online platforms (Australia, Brazil, China, Colombia, Ecuador, India, Malaysia, Mexico, South Africa, and the United States), creating an opportunity to discuss and standardise the data needed to monitor genetic diversity across the countries that represent the majority of the world’s biodiversity. Our results support the value and public benefit added by such open-access biodiversity information systems, which may be given for granted, but are only possible through deliberate, sustained national investment. Importantly, a smaller group of megadiverse countries maintains emerging or still-developing systems, or lack of a unified national infrastructure, which should be included in capacity building, knowledge exchanges and funding mobilization.

Finally, although the conservation of genetic diversity forms part of Mexico’s international commitments and is reflected in different components of the national legal framework, existing systems need to be strengthened and made more dynamic so that they can effectively support the long-term maintenance of genetic diversity (Hébert et al. 2026). Monitoring diversity should therefore extend beyond tracking its trajectories to safeguarding the evolutionary and biocultural processes through which genetic diversity is generated and maintained. In Mexico, these processes are threatened by different forms of agricultural management, infrastructure expansion, and economic development and land-use policies (Auliz-Ortiz et al. 2023). Their impacts could be mitigated through effective coordination between institutions responsible for biodiversity conservation and those whose decisions can affect evolutionary processes. Aligning public policies across institutions would help reduce contradictory actions and allow governmental decisions to be evaluated not only in terms of their economic and social outcomes (Moure et al. 2021), but also in terms of their consequences for genetic diversity and evolutionary potential. Ultimately, preventing threats before they affect populations remains one of the most effective conservation strategies, particularly because many such threats originate outside the environmental sector and beyond the scope of local management decisions (Phalan et al. 2017).

## CONCLUSIONS

This detailed analysis of data from Mexico’s first national assessment allowed us to clarify the importance of genetic diversity indicators within the country’s environmental information system and to demonstrate that they complement information already generated through NOM-059 assessments. Their incorporation reveals a dimension of vulnerability that may remain undetected when assessments focus solely on the current conservation status and characteristics of species, highlighting the need to consider the adaptive potential of populations and their capacity to persist across generations. Thus, these indicators are relevant not only for fulfilling international commitments, but also for strengthening biodiversity assessment and conservation at the national level. Their integration through communication, coordination, and continued monitoring among the actors involved could make an important contribution to conservation in a megadiverse country such as Mexico.

This first assessment also showed that Mexico has the scientific capacity, infrastructure, and information needed to begin implementing genetic diversity assessments at a broader scale. Rather than requiring parallel systems, integrating these assessments into existing national frameworks provides an opportunity to strengthen biodiversity monitoring and assessment by connecting genetic information with extinction-risk assessments and conservation actions.

Megadiverse countries concentrate most of the world’s biodiversity (Mittermeier et al. 2005), they encompass different levels of research infrastructure, and nearly all of them have been historically committed to the Convention of Biological Diversity, often setting precedents and leading new approaches. Therefore, although Mexico assessed relatively few species for its first report to the GBF, its experience is meaningful for global efforts. In particular, it shows that genetic diversity monitoring can build on information, expertise, and infrastructure that are already available, while gradually increasing the number of taxa assessed and improving coordination among researchers, government institutions, and local actors.

## ACKNOWLEDGMENTS

We are extremely thankful to the following colleagues, students and local experts who helped during discussions: Alejandra Barrios Flores, Angélica Cervantes, Priscila Duarte, Orlando Jael Espinosa Chávez, Ana Rebeca Flores Ramos, Roxana Abigail Flores Romero, Vania Jiménez Lobato, Patricia Koleff, Angélica Elizabeth Martínez Bauer, José Carlos Moreno Juárez, Adolfo Gerardo Navarro Siguenza, Annamaria Savarino-Drago, Romina Miranda, Emmanuel Texon Sánchez, Sofía Villarreal Ponce, Antonio González Rodríguez and Ricardo Gaytán Legaria. AW gratefully acknowledges PASPA-DGAPA-UNAM for supporting her sabbatical research stay at the Royal Botanic Gardens, Kew. MR-L, VA, and RA-G acknowledge SECIHTI postgraduate fellowships (CVU 2104410; CVU: 1033774, scholarship no. 762515; and 2103783, respectively), and support from their respective UNAM postgraduate programs: Ciencias Biológicas (MR-L, VA) and Ciencias de la Sostenibilidad (RA-G)

## AUTHORSHIP CREDIT

Conceptualization: AMY, AW, FAG. Software: AMY, VK, JdS. Methodology: AMY, AW, JdS, LL, SH. Formal analysis: AMY, AW. Data collection: LAA, JFO, MCLC, ALQ, SRB, EZ, DMRR, VA, JJSG, AMY, AW. Data curation: MCLC, RAG, MRL, SSA, ACA, MAOR, OOG, CB. Writing original draft: AMY, AW. Writing review and editing: everyone. Visualization AMY. Project admin AMY, AW.

## DATA AVAILABILITY STATEMENT

Clean, processed data and R code to conduct analyses and plots are available at the Zenodo repository XXXX (available upon acceptance).

